# Proline Stickland fermentation supports *C. difficile* spore maturation

**DOI:** 10.1101/2025.03.13.643084

**Authors:** Zavier A. Carter, Christopher E. O’Brien, Shonna M. McBride

## Abstract

*Clostridioides difficile* is an anaerobic pathogen that thrives in the metabolically diverse intestinal environment. *C. difficile* is readily transmitted due to its transformation into a dormant spore form that is highly resistant to heat and disinfectants. Nutrient limitation is a key driver of spore formation; however, few metabolites have been directly shown to influence the regulation of *C. difficile* sporulation. A distinct aspect of *C. difficile* biology is the fermentation of amino acids through Stickland metabolism pathways, which are critical sources of energy for this pathogen. We hypothesized that as a preferred energy source, the amino acid proline may serve as a signal that regulates the initiation of sporulation or the development of spores. Using mutants in the proline reductase gene, *prdA*, and the proline-dependent regulator, *prdR,* we examined the impact of proline on *C. difficile* physiology and differentiation. Our results demonstrate that proline reductase is important for the development of mature spores and that excess proline can repress *C. difficile* sporulation through PrdR regulation. Further, we discovered that the end product of proline reduction, 5-aminovalerate, can support the growth of *C. difficile* through an unidentified, PrdR-dependent mechanism.

**IMPORTANCE:** *C. difficile* is an anaerobic intestinal pathogen that disseminates in the environment as dormant, resilient spores. Nutrient limitation is known to stimulate spore production, but the contribution of specific nutrients to sporulation is poorly understood. In this study, we examined the contribution of proline and proline fermentation to spore formation. Our results demonstrate the effect of proline fermentation on spore quality and the importance of the proline reductase pathway on spore maturation.

## INTRODUCTION

*Clostridioides difficile* is an anaerobic gastrointestinal pathogen that causes recurrent and debilitating diarrheal disease. The two major features of *C. difficile* biology that contribute to pathogenesis are the production of destructive toxins (TcdA and TcdB) and the formation of resilient spores that facilitate the spread of the bacteria. The production of both *C. difficile* toxins and spores are stimulated by nutrient deprivation.(1–4) Although it is known that nutritional availability greatly influences *C. difficile* differentiation into spores, few specific nutrients or mechanisms that contribute to the regulation of spore formation have been identified or characterized.(5–8)

The main complication for understanding how nutrition influences *C. difficile* physiology is the breadth of metabolites and mechanisms used by the bacterium to generate energy. As an anaerobe, *C. difficile* derives much of its energy from glycolysis and fermentation. But in contrast to most pathogens, it employs ancient metabolic pathways for the fermentation of amino acids, ethanolamine degradation, electron bifurcation, and CO_2_ fixation, as well as fermentation of a variety of sugars to generate energy.(9–12) Of the substrates available to *C. difficile* in the intestine, proline was found to serve as a preferred substrate for energy generation through proline reductase, a Stickland fermentation pathway.(10, 13–15) The *C. difficile* proline Stickland pathway consists of six enzymes (PrdA,B,C,D,E,E2,F) that reduce D-proline to 5-aminovalerate. Transcription of the *prd* operon is induced by proline through the regulator, PrdR. Though proline is clearly important for *C. difficile* intestinal colonization and toxin production (13, 16), the impact of this nutrient on sporulation remains unknown.

In this study, we investigated the effects of proline and proline Stickland metabolism on *C. difficile* sporulation and spore maturation. We found that proline repressed sporulation of wild-type *C. difficile* and that proline fermentation was important for proper spore development and maturation. Our results demonstrate that proline reductase (PR) activity contributes to the activation of early sporulation factors, but the need for PR can be overcome with additional nutrient supplementation. We also observed that proline, rather than the end product 5-aminovalerate (AMV), elicits a response through the regulator PrdR. Further, we provide the first evidence that the proline fermentation product, AMV, supports *C. difficile* growth through an uncharacterized PrdR-dependent pathway.

## RESULTS

### Proline reductase supports proficient spore production

Proline is a preferred energy source for *C. difficile* and was previously shown to support optimal growth.(13, 14) The decreased availability of other favored nutrients, such as sugars and branched-chain amino acids (BCAA) are known to trigger sporulation, though few mechanisms that connect nutrient availability and spore formation have been characterized.(6–8, 17) To determine if proline availability or proline catabolism impact sporulation, we generated deletion mutants for a proline reductase gene, *prdA,* and the proline-responsive regulator, *prdR*, and examined their phenotypes during growth on sporulation agar (**Figure S1, Figure 1**).(**ref kip**) The *prdA* mutant produced fewer ethanol-resistant spores than the parent strain (6.5 +/- 1.9 vs 18.4 +/- 3.4), suggesting that proline reductase is important for spore formation. Conversely, the *prdR* mutant produced more spores on average than the wild-type (33.2 +/- 9.0 vs 18.4 +/- 3.4), but sporulation in this strain was variable and did not reach statistical significance. Restoration of *prdA* or *prdR* to the respective mutants complemented the sporulation phenotypes observed.

**Figure 1.**
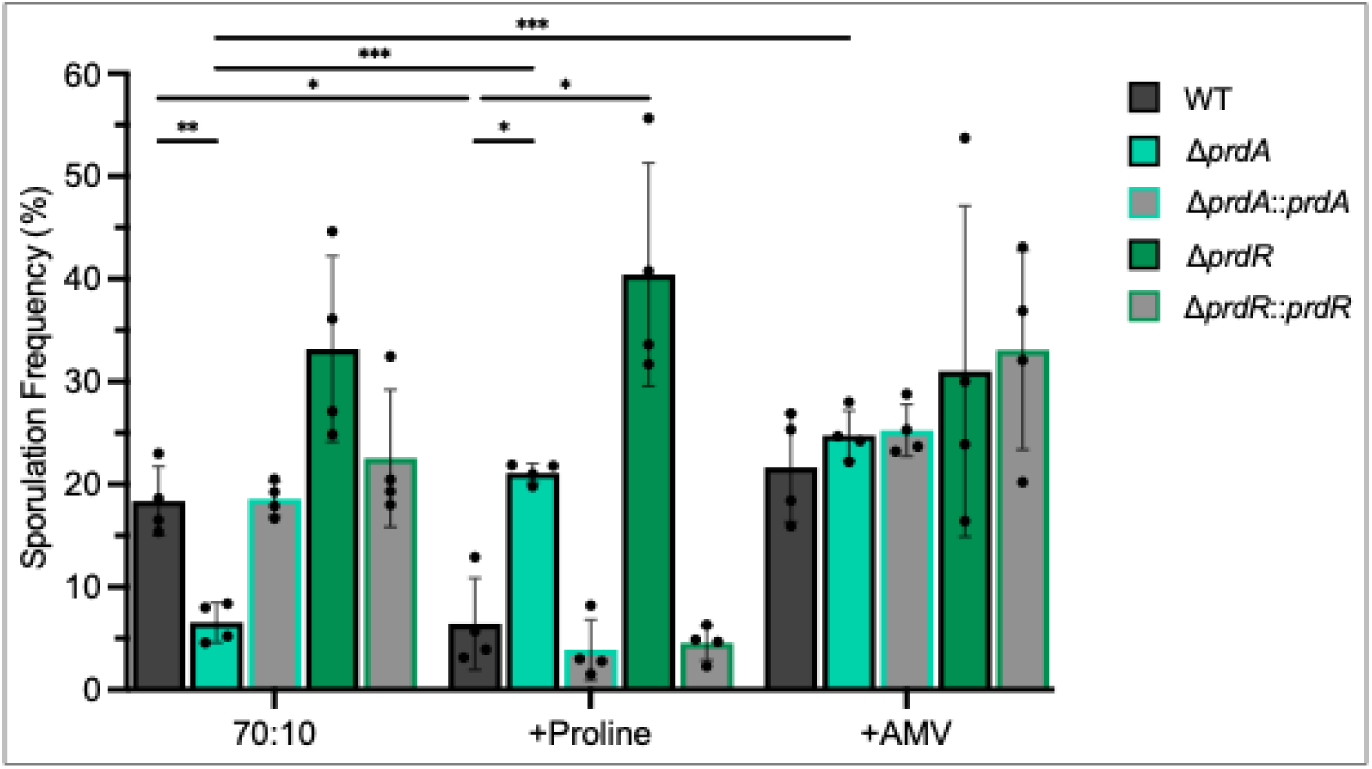
Proline reductase supports spore production, but is alleviated with nutrient supplementation. Ethanol-resistant spore formation for wild-type (630Δ*erm*)**, Δ***prdA* (MC2773), the Δ*prdA::prdA* complement (MC2774), Δ*prdR* (MC2337), and the Δ*prdR::prdR* complement (MC2668) grown on 70:10 sporulation agar +/- 30 mM proline or 5-aminovalerate. The means and individual values for four biological replicates are shown. Data were analyzed by two-way ANOVA with Dunnett’s multiple comparison test respective to the parent strain grown on 70:10. \**P* <0.05, \*\**P* <0.01, \*\*\**P* <0.001

To examine the impact of proline availability on sporulation, 30 mM proline was added to sporulation agar and strains were again assessed for spore production (**Figure 1**). The supplementation of proline to the medium reduced spore formation of the parent strain 2.8-fold, showing that proline can repress sporulation, similar to the repressive effects observed with sugars.(6) However, proline had the opposite effect on the *prdA* mutant, resulting in a 3.2-fold increase in sporulation for this strain. The increased sporulation of the *prdA* mutant with proline suggests that proline catabolism is not the only mechanism by which proline affects sporulation, but also likely occurs through the conversion of proline to alternative catabolites. Unlike the *prdA* mutant or wild-type strain, the *prdR* mutant sporulation frequency remained elevated with proline supplementation, indicating that the sporulation response to proline involves PrdR regulation.

Although proline catabolism reduces sporulation of wild-type *C. difficile*, the effector required for this response is not known. Regulation can occur in direct response to a catabolite or from response to a metabolic end product.(1, 6) Thus, the response to proline may be direct or a response to the metabolic product of proline reductase, 5- aminovalerate. To test whether proline and proline reductase sporulation effects are due to feedback from proline metabolism, we supplemented the sporulation medium with 30 mM 5-aminovalerate (AMV, **Figure 1**), the product of proline fermentation. AMV supplementation had no impact on sporulation of the parent strain, indicating that the response to proline is direct, rather than feedback from proline catabolism.

Unexpectedly, the addition of AMV increased sporulation of the *prdA* mutant, leading to 3.8-fold greater spore formation. The *prdR* mutant again demonstrated elevated spore production with no apparent response to AMV. These results suggest that *C. difficile* can utilize AMV independently of proline reductase, but that AMV does not directly elicit a regulatory response that impacts sporulation.

### Proline and aminovalerate are catabolized by *C. difficile* through independent PrdR-regulated mechanisms

The AMV-dependent increase in sporulation observed for the *prdA* mutant was surprising, as it suggested that the *prdA* mutant catabolized AMV. To understand how AMV impacts *C. difficile,* we next investigated the effect of AMV on growth of wild-type (630Δ*erm*), the *prdA*, and *prdR* mutant. *C. difficile* cultures were grown in TY broth alone or with the addition of 30 mM proline or AMV (**Figure 2A,B**). As previously observed, proline enhanced the growth of the wild-type strain, but had no significant effects on the growth of the *prdA* or *prdR* mutants.(4, 13, 14) AMV also improved the growth of the parent strain, but had the greatest effect on growth for the *prdA* mutant, and no impact on growth of the *prdR* strain (**Figure 2**). These results demonstrate that AMV is utilized by *C. difficile* to support growth, which was not previously known. Further, the ability of the *prdA* mutant to catabolize AMV demonstrates that AMV is not utilized by a reversal of the proline reductase pathway, as AMV utilization occurs in the absence of proline reductase activity. Therefore, AMV catabolism occurs via a separate, unknown pathway.

**Figure 2.**
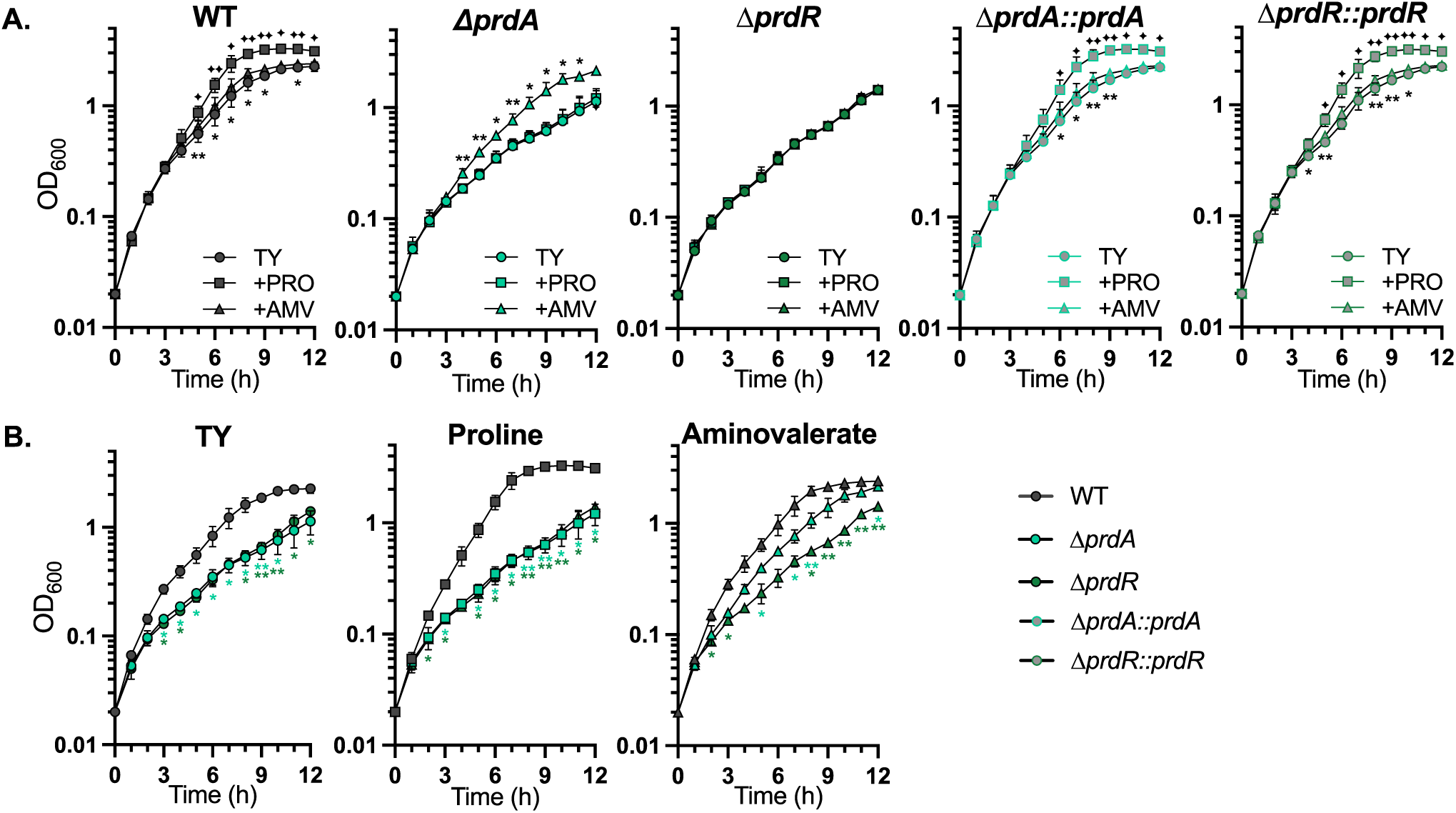
PrdR independently regulates growth in proline and aminovalerate. **A)** Growth of wild-type (WT, 630Δ*erm*), **Δ***prdA* (MC2773), the Δ*prdA::prdA* complement (MC2774), Δ*prdR* (MC2337), and the Δ*prdR::prdR* complement (MC2668) grown in TY with and without 30 mM proline or 5- aminovalerate. **B)** Condition-centered display of growth for WT, Δ*prdA*, and Δ*prdR* mutant in TY with and without 30 mM proline or 5-aminovalerate. The means and SD of three independent experiments are shown. A one-way ANOVA with Dunnett’s multiple comparison test was used to compare the mutants to the parent strain at each timepoint and condition. A two-tailed Student’s *t* test was used to assess complemented strains with the relevant mutant. A) *, ✦, *P* <0.05 for aminovalerate or proline; B) *color indicates strain comparison; two symbols represents *P* <0.01.

The observation that growth of the *prdR* regulator mutant was unchanged with proline confirms prior work showing that PrdR regulates proline-dependent growth.(13, 14) In addition, we show that AMV supplementation has no effect on *prdR* mutant growth, demonstrating that PrdR controls AMV-dependent growth (**Figure 2A,B**). All growth defects observed with the *prdA* and *prdR* mutants were restored with complementation of the respective genes (**Figure 2A**).

### Proline reductase facilitates the maturation of spores

In addition to the evaluation of spore ethanol-resistance, we examined the *prdA* and *prdR* mutant spore formation by phase-contrast microscopy (**Figure 3A**). The *prdR* mutant had variable production of mature or “phase-bright” spores, similar to results observed for ethanol resistance (**Figure 1**). In contrast, the *prdA* mutant produced fewer phase-bright spores than the parent strain when grown on sporulation agar. However, the *prdA* mutant also made more immature or “phase-dark” spores than other strains, as shown quantified in **Figure 3B**. These data reveal that the *prdA* mutant is able to initiate sporulation, but the population forms mature spores at a low frequency (**Figure 1**).

**Figure 3.**
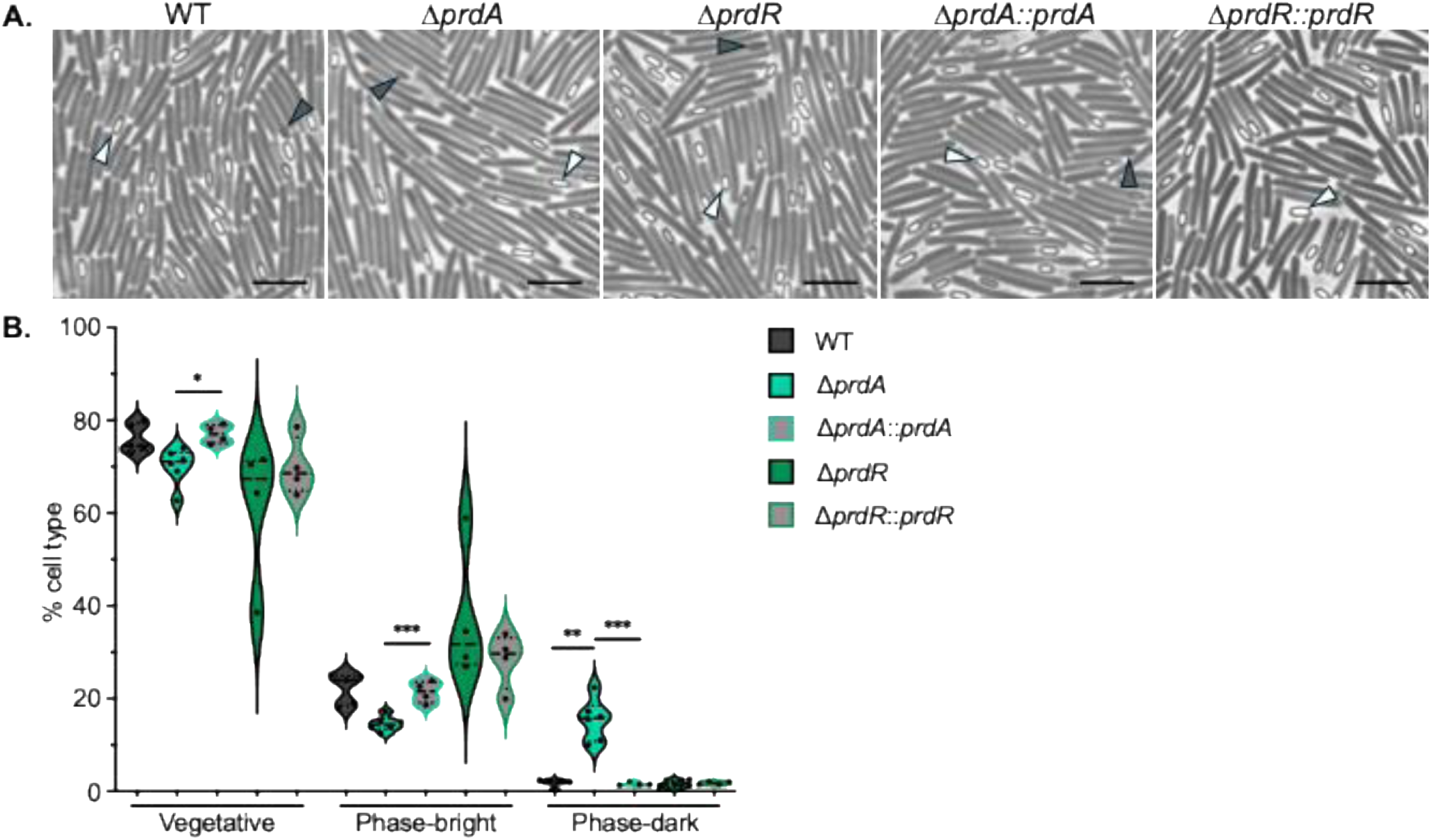
Proline reductase facilitates complete spore formation. **A)** Phase- contrast micrographs of wild-type (630Δ*erm*)**, Δ***prdA* (MC2773), the Δ*prdA::prdA* complement (MC2774), Δ*prdR* (MC2337), and the Δ*prdR::prdR* complement (MC2668) grown on 70:10 sporulation agar for 24 h. White arrowheads indicate phase bright spores and dark arrowheads indicate phase dark spores. Scale bar = 5 µm. **B)** Incidence of cell morphologies following growth on sporulation agar for the strains above. Data for approximately 1000 cells were assessed for a minimum of four biological replicates. Median values are indicated by a heavy black line; quartiles are represented as dotted lines within violin plots. Statistical analyses: one-way ANOVA with Dunnett’s multiple comparisons test was performed for mutants relative to WT; unpaired Student’s *t* tests comparing the mutants to their respective complemented strain. \**P* <0.05, \*\**P* <0.01, \*\*\**P* <0.001

### Proline consumption during sporulation supports efficient spore germination

The decreased sporulation of the *prdA* mutant led us to hypothesize that the resulting spores produced may be underdeveloped. An important characteristic of mature spores is the development of sensitivity to germinants and the mechanisms necessary for germination.(18) To test the impact of proline catabolism on spore development, we assessed the *prdA* and *prdR* mutants for germination using purified phase-bright spores. As shown in **Figure 4**, germination was significantly delayed for the *prdA* mutant spores, while the *prdR* mutant displayed a germination rate similar to the parent strain. Together, the slow germination of the *prdA* mutant and its reduced production of phase-bright spores are indicative of defective spore development with the loss of proline reductase activity.

**Figure 4.**
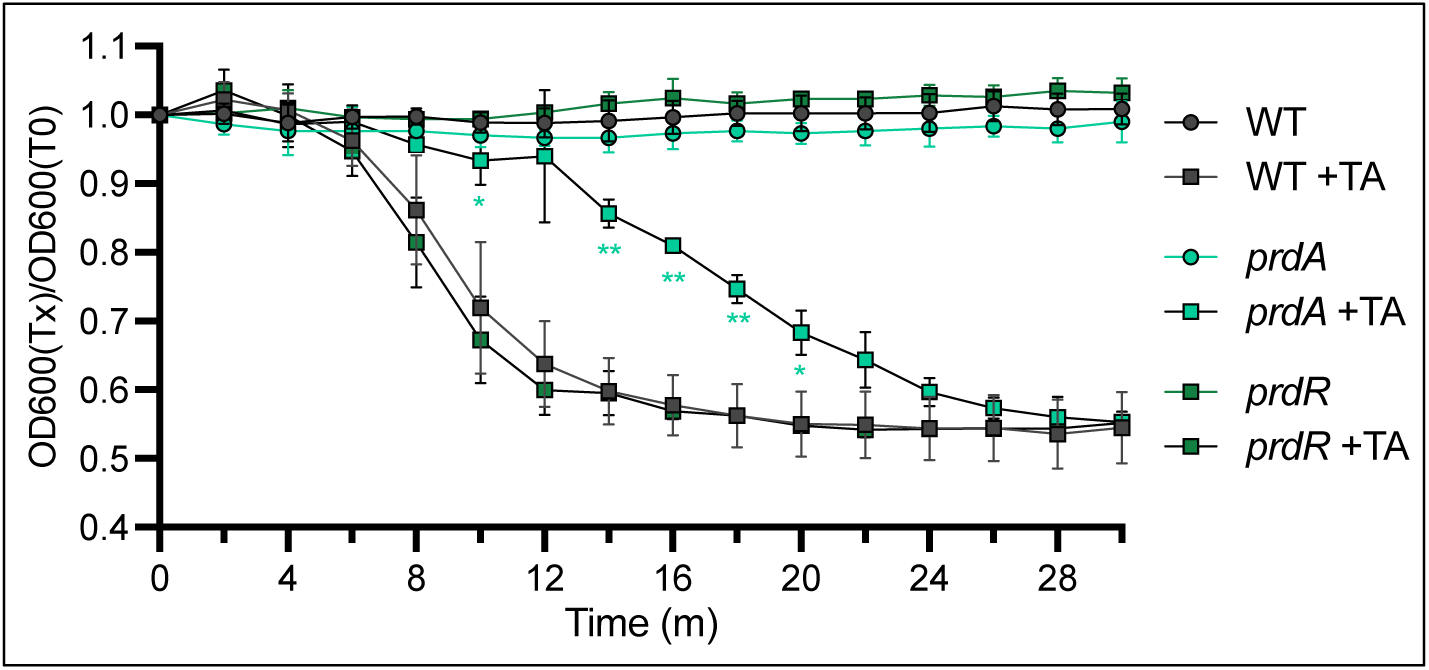
Proline utilization during sporulation supports efficient spore germination. Purified *C. difficile* spores from wildtype (WT, 630Δ*erm*)**, Δ***prdA* (MC2773), and Δ*prdR* (MC2337), were assayed for germination in BHIS with or without the germinant taurocholate. Optical densities of spore samples were assessed over time and the ratio of the optical densities for each timepoint relative to the initial optical density are shown. The means are standard deviation for at least three independent experiments are shown. A two-way ANOVA with Tukey’s multiple comparison test was used to compare the mutants to the parent strain for each timepoint and condition. \**P* <0.05, \*\**P* <0.01.

### Loss of proline reductase activity impairs expression of late-stage sporulation genes

Successful spore formation requires the sequential activation of the four sporulation-specific sigma factors: SigF, SigE, SigG, and SigK.(19–21) The sporulation sigma factors are transcribed and held inactive until an activation signal is coordinated by the neighboring compartment (mother cell or forespore). Once activated, each sigma factor controls the compartment-specific expression of genes required for a phase of spore development.(20, 21) To understand how proline catabolism impacts the progression of sporulation, we examined the temporal expression of sporulation sigma factors and genes that are transcribed by each sigma factor during the phases of development (**Figure 5; Figure S2**). First, transcript levels of the early forespore sigma factor, *sigF*, and the SigF-dependent gene, *gpr*, were unaffected in the *prdA* or *prdR* mutants, relative to the parent strain. Transcript levels of the initial mother cell sigma factor *sigE* were also unaffected in the mutants, but expression of the SigE-dependent transcript, *spoIID,* was modestly reduced in both the *prdA* (0.5 ± 0.1) and *prdR* mutant (0.6 ± 0.1), suggesting decreased activation of SigE in the mother cell. Transcript levels of *sigG,* encoding the secondary forespore sigma factor, were unchanged in the *prdA* and *prdR* mutants. However, the delay in SigE activation observed for both mutants carried into subsequent phases of sporulation, leading to reduced activation of SigG (*sspA* expression), with the *prdA* mutant displaying the greatest deficiency (0.18 ± 0.1). For the *prdA* mutant, these delays culminated in significant decreases in transcription of the late-stage mother cell sigma factor, SigK, (0.15 ± 0.06) and the SigK-dependent gene, *cotA* (0.07 ± 0.05). In summary, the *prdA* mutant has reduced activation of early sporulation factors that compound through the regulatory cascade and lead to further deficits at later stages which results in a low frequency of mature spores. Hence, the energy generated by the reduction of proline is important for the initiation of sporulation and the ability to orchestrate spore development.

**Figure 5.**
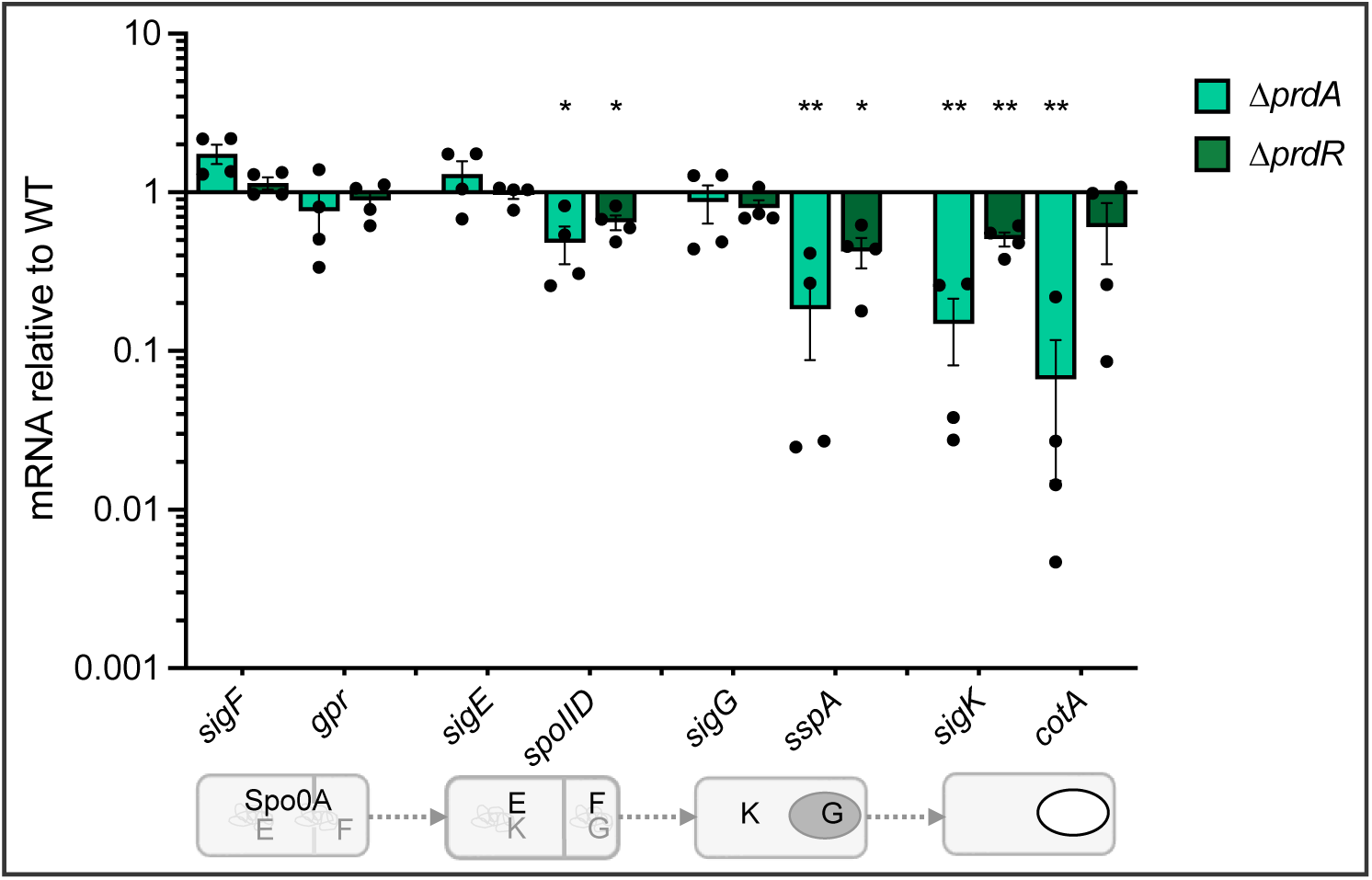
Loss of proline reductase impairs expression of late-stage sporulation genes. Quantitative reverse transcription PCR analysis of sporulation-specific genes temporally expressed during spore development for **Δ***prdA* (MC2773), and the Δ*prdR* mutant (MC2337) grown on 70:10 sporulation agar for 12 h, relative to the parent strain 630Δ*erm*. The bottom panel depicts the sequential expression (grey) and activation (black) of the sporulation sigma factors during spore development. The means, individual values, and standard error of the mean are shown for four biological replicates. Data were analyzed by two-way ANOVA with Dunnett’s multiple comparison test respective to the parent strain grown on 70:10. \**P* <0.05, \*\**P* <0.01

### Proline catabolism broadly impacts nutritional regulation

Proline availability is sensed by the PrdR regulator, which drives expression of the proline reductase (*prd*) operon. The reduction of proline results in the generation of NAD^+^, which in turn is sensed by the redox responsive regulator, Rex.(22) Hence, the PrdR and Rex regulons are interrelated. Rex regulates the transcription of many metabolic genes and operons including glycine reductase, succinate utilization, butyrate production, ethanolamine degradation, and cobalamin biosynthesis.(22) We considered that the loss of proline reductase may create a redox imbalance that disrupts sporulation progression. To test this, we examined the transcription of the Rex and PrdR-regulated genes, *grdE* and *CD2344*, which are associated with proline availability and redox response (**Figure S3**). No significant changes were observed for either transcript in the *prdA* or *prdR* mutant, suggesting that Rex and PrdR-dependent regulation are not impacted by the loss of proline reductase.

Since no significant Rex or PrdR regulation was apparent, we hypothesized that the *prdA* mutant may have changes in the regulation of other nutritional genes that compensate for the loss of proline reductase activity. To this end, we investigated nutritional transcripts that are controlled the carbon catabolite regulator, CcpA, and the BCAA-GTP responsive regulator, CodY (**Figure 6**).(6, 8, 13, 14, 17, 22) Both CcpA and CodY suppress sporulation and toxin expression in *C. difficile*, as well as regulate hundreds of genes involved in nutrient acquisition.(2, 6–8) Modest reductions were found in the transcript levels of the CcpA-repressed gene, *CD0314* (0.46 mean ± 0.20 SEM), and the co-repressed oligopeptide permease gene, *oppB* (0.43 mean ± 0.11 SEM). Reduced transcription of these genes suggests that the *prdA* mutant incompletely responds to the loss of proline as an energy source to increase nutrient acquisition.

**Figure 6.**
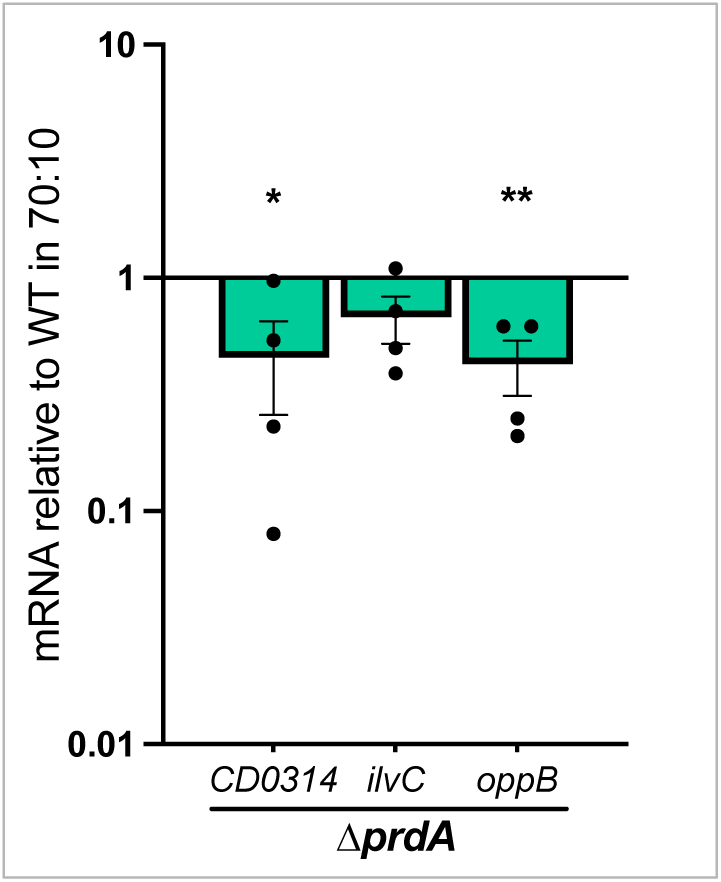
Proline utilization broadly impacts nutritional regulation. Quantitative reverse transcription PCR analysis of nutritionally-regulated genes *CD0314* (CcpA-dependent), *ilvC* (CodY-dependent), and *oppB* (CodY and CcpA- dependent) for the **Δ***prdA* mutant (MC2773) grown on 70:10 sporulation agar for 12 h, relative to the parent strain 630Δ*erm*. The means, individual values, and standard deviation for four biological replicates are shown. Unpaired Student’s *t-* tests comparing the mutants to their respective complemented strain. \**P* <0.05, \*\**P* <0.01.

## DISCUSSION

Work in other spore-forming species of *Bacillus* and *Clostridium* have demonstrated that nutrition plays a major role in the rate, frequency, and hardiness of spores, though the specific nutrients that impact this process varies by species.(23–27) Our objective was to determine what role proline and proline fermentation have in the regulation of *C. difficile* sporulation and the development of spores. Our results demonstrate that the availability of abundant proline acts as a signal that can impede spore formation, as observed with proline supplementation to sporulation medium (**Figure 1**). Prior work with other preferred carbon sources, such as glucose, fructose, and BCAA showed that abundance of favored nutrients inhibits spore formation in *C. difficile* by signaling through global regulators (CcpA and CodY).(6, 7) Similarly, we observed that excess proline leads to repression of sporulation through the sigma-54 dependent regulator, PrdR. PrdR was previously found to regulate more than 50 genes involved in the uptake and metabolism of nutrients for energy and redox balance, though no sporulation genes were noted among PrdR-regulated transcripts.(22) However, regulation by PrdR is closely tied to the activity of the redox regulator, Rex, and it is not clear whether the proline-dependent sporulation phenotypes observed are mediated by PrdR directly or by Rex.

We also showed that proline reduction serves as an important energy source during the initiation of *C. difficile* sporulation, and allows for the activation of early sporulation factors that are required for spore development (**Figure 1, 3, 5**). In the absence of proline reductase activity, the majority of cells failed to form mature spores, resulting in poor development of ethanol resistance and delayed germination (**Figure 3, 4**). In a previous study investigating the effects of glycine Stickland fermentation, we also observed a decrease in spore maturation in a glycine reductase (*grd*) mutant.(5) However, the spores that were produced by the *grd* mutant displayed more rapid germination than the parent strain.(5) Unlike the *prd* mutant, which demonstrated stalling of early (SigE-dependent) sporulation gene expression, the *grd* mutant maintained wild-type levels of sporulation transcripts up to the final sporulation sigma factor, *sigK.* These differences in the phenotypes of *grd* and *prd* mutants suggest that these energy-generating pathways have distinct roles in supporting the developing spore, with proline reductase playing a greater role during early stages and glycine reductase at the final stages.

An unexpected result from this study was the observation that *C. difficile* can metabolize the proline reductase end product, 5-aminovalerate (AMV). While it has been known for more than 100 years that proline can be degraded to AMV, this is the first report to our knowledge of AMV as a substrate for energy generation.(28–30) Given that the growth of the *prdA* mutant is enhanced by supplementation with AMV (**Figure 2**), we can infer that AMV utilization does not occur through a reverse reaction via the proline reductase pathway. The identity of the mechanism for AMV metabolism is not known, but the loss of AMV-dependent growth enhancement in the *prdR* mutant suggests that AMV is utilized through a PrdR-dependent pathway. Of the published PrdR-dependent genes, we did not find metabolic factors that were clear candidates for AMV degradation, thus further investigation is needed to identify the factors involved.(22) In addition, a greater understanding of the ability of proline and proline reductase activity to manipulate spore initiation and maturation may be exploited to decrease spore formation, and thereby reduce transmission.

## MATERIALS AND METHODS

### Strain and plasmid construction

The bacterial strains and plasmids used for this study are found in **Table 1**. Primers for *C. difficile* were designed using the 630 strain genome (GenBank accession no. NC_009089.1) and listed in **Table 2**. All constructs were sequenced by Plasmidsaurus. *C. difficile* mutants were constructed using pseudo-suicide allele-coupled exchange as previously described. (31, 32)

**Table 1.**
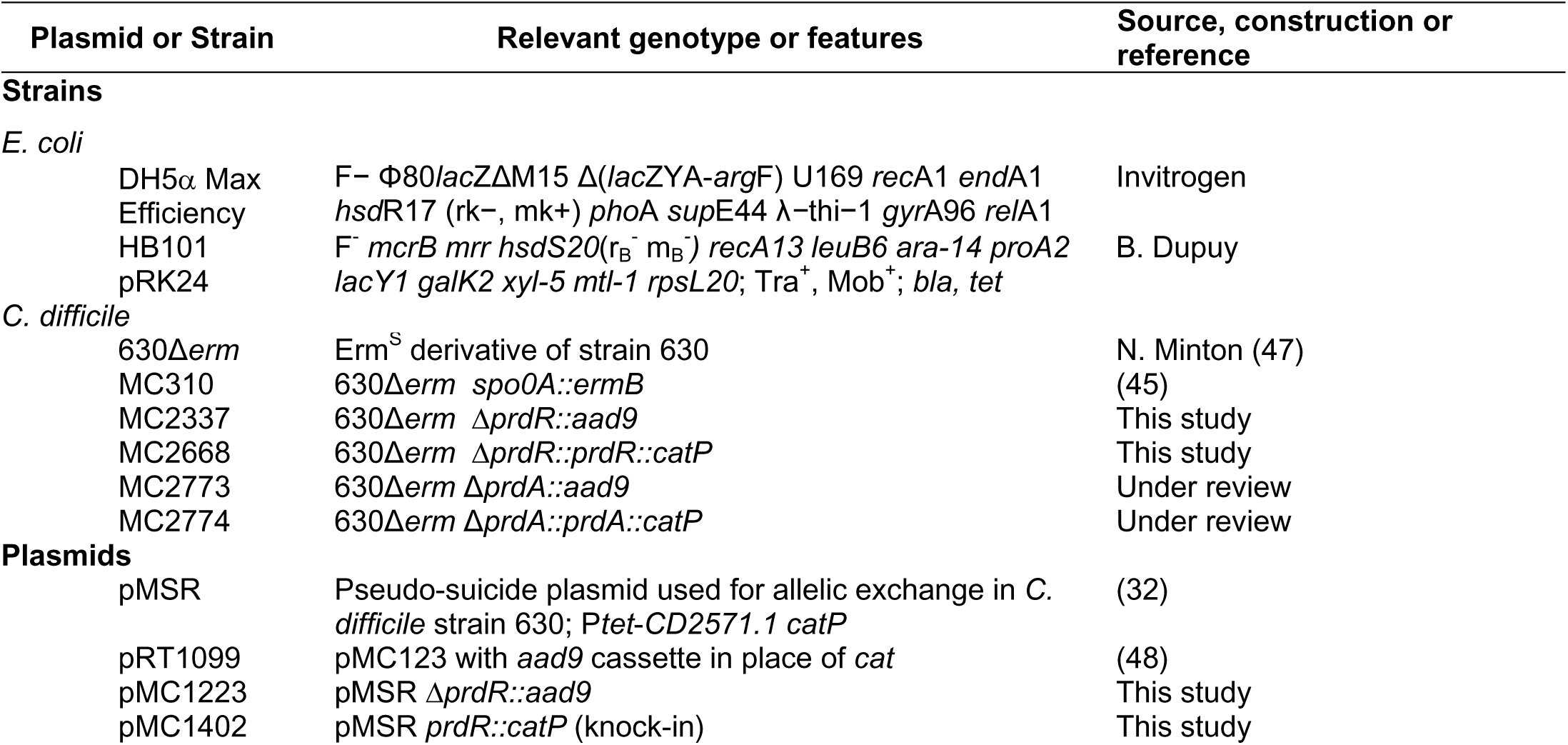
Bacterial Strains and plasmids.

**Table 2.**
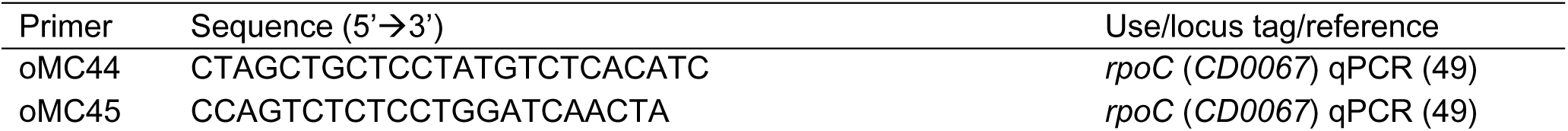
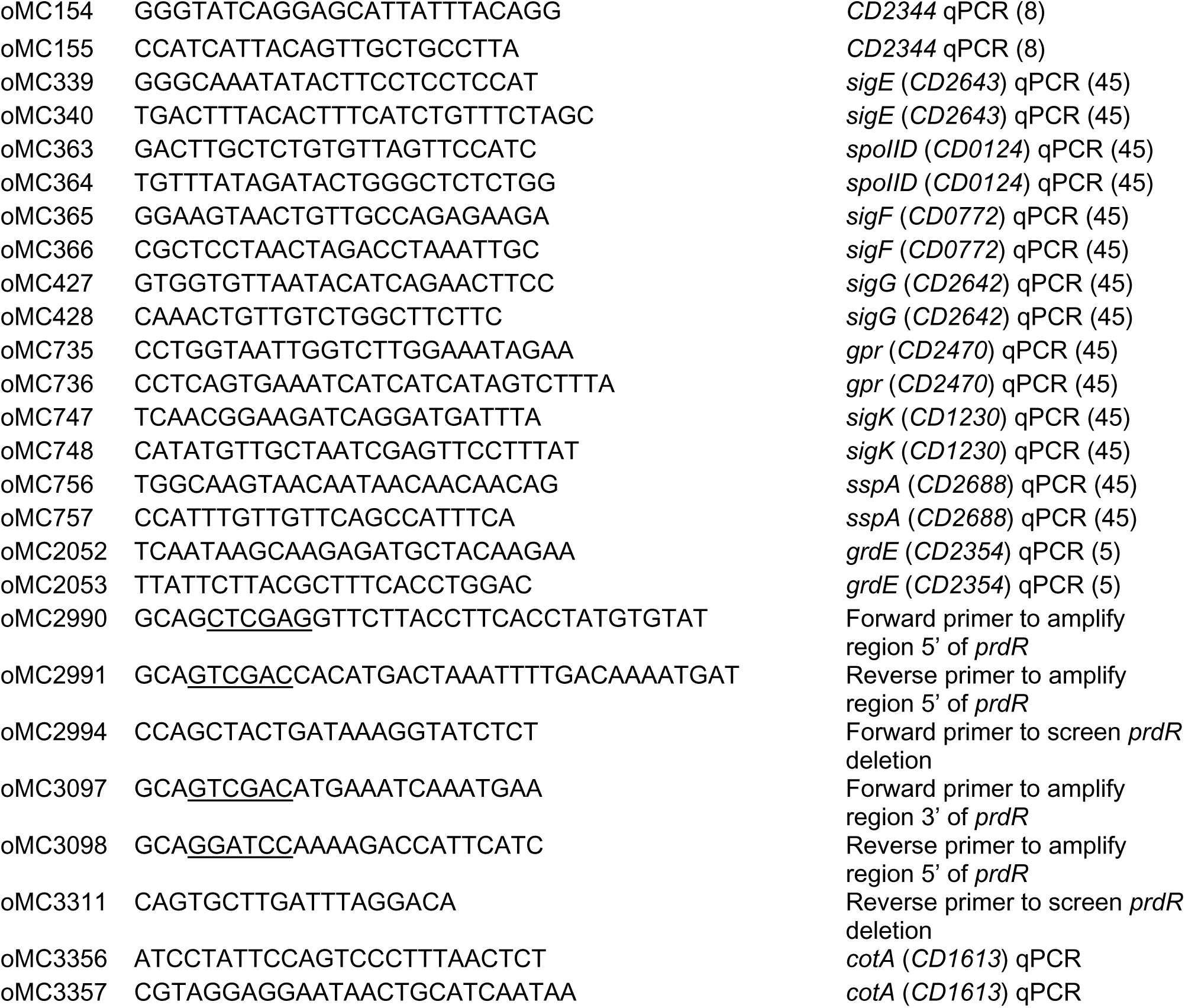
Oligonucleotides.

### Whole Genome Sequencing

*C. difficile* genomic DNA was extracted and sent for whole genome sequencing as previously described.(33) Genomic library preparation and sequencing for the *prdA* and *prdR* deletion mutants were performed by SeqCenter (Pittsburgh, PA) on an Illumina NextSeq2000 platform (Edwards and McBride, 2023). Geneious Prime (v2023.1.2 and v.2021.2.2) was used to trim reads with the BBDuk plug-in and mapped them to the *C. difficile* reference genome NC_009089. The Bowtie2 plugin was used to find SNPs or InDels under default settings with a minimum variant frequency of 0.9 (34). Genome sequence files for the *prdA* mutant were deposited into NCBI Sequence Read Archive (SRA) Bioproject PRJNA1085672. The genome sequence files for the *prdR* mutant were deposited into NCBI SRA Bioproject PRJNA1231098.

### Bacterial growth assays and conditions

*Escherichia coli* strains were grown aerobically in LB medium (Lennox) at 37°C with 100 μg/mL spectinomycin (Chem-Impex) or 20 μg/mL chloramphenicol (Sigma-Aldrich) added for plasmid maintenance. Cultivation of *Clostridioides difficile* strains was performed at 37°C in a Coy anaerobic chamber with an atmosphere comprised of 85% nitrogen, 10% hydrogen, and 5% CO_2,_ as previously described.(35) Each *C. difficile* strain was grown in TY (36) or BHIS broth supplemented with 0.1% taurocholic acid (TA, Sigma-Aldrich) and 0.2% fructose (D-fructose, Fisher chemical) to induce germination and prevent sporulation, respectively, prior to growth assays.(37) For *C. difficile* transformation, 5 μg/mL of thiamphenicol (Sigma-Aldrich) and 500 μg/mL of spectinomycin was used for plasmid maintenance or integrant selection, as necessary. For counterselection against *E. coli*, 0.1% taurocholic acid and 100 μg/mL of kanamycin (Sigma-Aldrich) were used after conjugation. Stocks of 1 M L-proline (Sigma-Aldrich) and 1 M 5-aminovalerate (Tokyo Chemical Industry) were made with dH_2_O for use in growth both curves and spore assays For growth curves, active starter cultures were grown to an OD_600_=0.5, then 2 mL of culture was used to inoculate 48 mL of TY media +/- each supplement and monitored over 12 h.

### Sporulation assays

Sporulation assays were performed as previously described, with minor modifications to the sporulation medium.(38, 39) The original sporulation medium, 70:30, consisted of 70% SMC and 30% BHIS.(40) Due to increased potency of Difco BHI lots, the sporulation agar was adjusted to 10% BHIS, resulting in 70:10 medium. Active *C. difficile* cultures were applied to 70:10 agar as and incubated for 24 h prior to ethanol-resistance assays and plating for CFU, as previously described.(39) Sporulation frequency was calculated as the ratio of ethanol-resistant spores divided by the total number of cells/ml. A sporulation defective *spo0A* mutant was used as a negative control to ensure vegetative cell death from ethanol in all assays.

### Spore germination assays

*C. difficile* was incubated for greater than 48 h on 70:10 sporulation agar to induce spore formation and purified, as previously described.(41–43) Briefly, cells were scraped from sporulation agar and resuspended in spore buffer (1x PBS + 1% BSA), followed by centrifugation over a 12 ml 50% sucrose bed volume until samples reached greater than 95% purity of phase-bright spores. Spores were washed and resuspended in spore buffer and applied to germination assays at a starting density of ∼ OD_600_ 0.3. Germination frequency was calculated as the loss of OD_600_ upon addition of a 5 mM final concentration of the germinant taurocholate, as previously described.(43) Germination assays were performed in technical duplicate for three biological replicates and analyzed by one-way ANOVA for each 2 min timepoint over 30 min.

### Quantitative reverse transcription PCR analysis (qRT-PCR)

*C. difficile* strains were grown on 70:10 sporulation agar with or without supplementation, as described above for sporulation assays. Following 12 h of growth, cells were scraped and resuspended to a 1:1:2 Ethanol-Acetone-water solution and stored at -70°C. RNA was extracted as previously described, treated with DNase I (Ambion), and cDNA synthesized using random hexamers (Bioline).(44, 45) qRT-PCR was performed on a Roche LightCycler 96 instrument from 50 ng cDNA in technical triplicate using Bioline SensiFast SYBR & Fluorescein mix with the primers listed in **Table 2**. Expression was normalized to the internal control transcript, *rpoC* and analyzed following the ΔΔC_t_ method for relative quantification.(46) GraphPad Prism v10.4.1 was used for all statistical analyses, as indicated in figure legends.

## ACKNOWLEDGEMENTS

The authors would like to thank Adrianne Edwards and members of the McBride lab for useful feedback and suggestions during the course of this work. This work was supported by the U.S. National Institutes of Health through research grants AI156052 and AI116933 to S.M.M. The content of this manuscript is solely the responsibility of the authors and does not necessarily reflect the official views of the National Institutes of Health.

